# Plipastatin is a shared good by *Bacillus subtilis* during combating *Fusarium* spp

**DOI:** 10.1101/2024.12.07.627343

**Authors:** Rune Overlund Stannius, Ákos T. Kovács

## Abstract

*Bacillus subtilis* a Gram-positive soil dwelling bacterium known for its wide range of bioactive secondary metabolites. The lipopeptide plipastatin produced by most *B. subtilis* isolates have been shown to exhibit potent anti-fungal activity against plant pathogenic fungi. While the effect of these anti-fungal compounds are well studied in the context of biocontrol, much less is known of their role in the environment, which also harbor non-producing strains of these compounds. *Fusarium* species produce multiple antibacterial compounds resulting in dysbiosis of the plant-associated microbiome and inhibiting plant beneficial bacteria like *B. subtilis*. While plipastatin is expected to be important for survival of *B. subtilis*, not all isolates carry the biosynthetic gene cluster for plipastatin suggesting that the protective effect of plipastatin might be shared. In this study, we investigated the protective effect of plipastatin against *Fusarium oxysporum* in a co-culture using a producer and a non-producer isolate of plipastatin. We tested the survival of single and co-cultured strains under *Fusarium* challenge in liquid media and solid agar plates to dissect the influence of spatial structure. Our results highlights that plipastatin protects the non-producer strain in a density dependent manner.

## Introduction

Microbial interactions are the drivers of host health and disease, from individual microbe-microbe interactions to the whole network that make up the host microbiome. The many interactions that take place in the microbiome around and in a plant is one of the main factors contributing to the host fitness (Mendes, Garbeva and Raaijmakers 2013; Compant *et al*. 2019). Numerous bacteria have been shown to possess plant beneficial effects, by promotion of growth, nutrient utilization, or protection against plant pathogens either by induction of systemic resistance or direct inhibition of the plant disease-causing agent (Lugtenberg and Kamilova 2009; Raaijmakers *et al*. 2009; Blake, Christensen and Kovács 2021).

*Fusarium oxysporum* is a filamentous fungus and causative agent of Fusarium wilt in several economically important crops including banana, tobacco, and tomato (Summerell 2019). The disease is cause of significant yield losses due to clogging the vascular vessels of the plant leading to wilting of the leaves and death of the plant. Among fungal plant pathogens, *F. oxysporum* is placed as the fifth most scientifically and economically important fungal pathogen due to the breadth of its host range and severe impact on several large sectors of the agricultural market (DEAN *et al*. 2012). Historically, chemical fungicides have been employed in combating FHB, which has led to widespread ecological consequences and risk of resistance developing in *F. oxysporum* (Bawa 2016; Hudson *et al*. 2021; El-Aswad *et al*. 2023) underlining the need for more environmentally friendly biocontrol agents.

*Bacillus subtilis* group species are known to produce a suite of bioactive secondary metabolites, including lipopeptides such as surfactin, fengycins, and iturin (Ongena and Jacques 2008), including plipastatin that belongs to fengycins (Honma *et al*. 2012; Gong *et al*. 2015), which together with surfactin elicits systemic resistance in plants (Ongena *et al*. 2007; Farace *et al*. 2015). Plipastatin variants exhibit species-specific inhibition of *Fusarium*, while some plipastatin was reported to only affect *Fusarium graminearum* and not *F. oxysporum* (Zhao *et al*. 2014; Gong *et al*. 2015), others exhibit activity against both (Kiesewalter *et al*. 2021) making *Bacillus* a possible candidate for future biocontrol of the fungal pathogen. *Fusarium* produces several antibacterial compounds, which negatively affect the plant associated microbial community (Sondergaard *et al*. 2016; Venkatesh and Keller 2019), leading to dysbiosis and death of plant beneficial bacteria underlining the importance of effective countermeasures. Therefore, it is also expected that production of *Fusarium*-inhibiting metabolites, like plipastatin, could benefit the bacterial community by combating the invading fungi.

Interestingly, natural isolates of *Bacillus subtilis* were identified to possess or lack the production of plipastatin, including isolates that originate from the same soil samples (Kiesewalter *et al*. 2021), which raises the question whether the protective effect of plipastatin is shared with the species and broadly the microbial community. While sharing of plipastatin could protect the community, it might also suffer from cheating by other non-producing *Bacilli,* which could potentially out-compete the producer strain due to fitness costs and this could lead to reduced level of producing strains and a loss of plipastatin protection in the community. The lack of plipastatin and fengycin production has been broadly proposed in wide variety of isolates of *B. subtilis* and *Bacillus velezensis*, respectively, as either frame shift mutations or partial deletion have been identified in the respective biosynthetic gene clusters of these secondary metabolites (Kiesewalter *et al*. 2021; Steinke *et al*. 2021).

In this study, we evaluate the role of plipastatin in a co-culture system of two *B. subtilis* isolates when challenged by *F. oxysporum.* The two *B. subtilis* isolates originate from the same sample site, but one, MB9_B1, is a producer of plipastatin and thus able to inhibit *Fusarium* while the other isolate, MB9_B6, is unable to produce this secondary metabolite (Kiesewalter *et al*. 2021). In our co-culture setup, we demonstrate that the plipastatin producer can provide protection to the non-producer in a density dependent manner.

## Materials and methods

### Strains, culturing, and genetic modification

Strains used in this study can be found in Table 1. *B. subtilis* was routinely cultured in lysogeny broth (LB, Lenox, Carl Roth) for overnight cultures while *F. oxysporum* was cultured in PDB (Potato dextrose broth, Carl Roth). Strains were genetically engineered using the protocol below which is based on (Kunst and Rapoport 1995).

**Table 1.** Strains used in the study. *gfp: gfpmut2*, spec^R^: spectinomycin resistance, tet^R^: tetracycline resistance.

| Strain | Characteristics | Reference |
| --- | --- | --- |
| *Bacillus subtilis* MB9_B1 | natural isolate, plipastatin producer | (Thérien *et al.* 2020) |
| *Bacillus subtilis* MB9_B6 | natural isolate, plipastatin non-producer | (Kiesewalter *et al.* 2021) |
| *Bacillus subtilis* MB9_B1 mKate2 | MB9_B1 transformed with gDNA from TB501 *amyE::P<sub>hyperspank</sub>-mKate2* (*spec<sup>R</sup>*) (Dragoš *et al.* 2018) | This study |
| *Bacillus subtilis* MB9_B1 mKate2 Δ*ppsC* | MB9_B1 mKate2 transformed with gDNA from DS4114 Δ*ppsC::tet<sup>R</sup>* (Müller *et al.* 2014) | This study |
| *Bacillus subtilis* MB9_B6 GFP | MB9_B6 transformed with gDNA from TB500 *amyE::P<sub>hyperspank</sub>-gfp* (*spec<sup>R</sup>*) (Mhatre *et al.* 2017). | This study |
| *Fusarium oxysporum* IBT 40872 | From surface of an equipment in a Danish factory, 2005, by Anne Svendsen | IBT Culture Collection of Fungi, DTU |

Bacterial inoculants were prepared from 1 mL overnight cultures, which were spun down and resuspended in 100 µL MQ water. 10 µL of the inoculum was transferred into 2 ml of competence medium (80 mmol/L K_2_HPO_4_, 38.2 mmol/L KH_2_PO_4_, 20 g/L glucose, 3 mmol/L Na_3_-citrate, 45 µmol/L ferric NH_4_-citrate, 1 g/L casein hydrolysate, 2 g/L K-glutamate, 0.335 µmol/L MgSO_4_·7H_2_O) and incubated at 37 °C with shaking for 3.5 hours. Donor DNA was prepared from 1 ml of an overnight culture of the donor strain using Bacterial and Yeast Genomic DNA Purification Kit from EURx, yielding a typical DNA concentration in the range of 50 – 150 ng/µL. After the initial growth phase, 2 µL donor DNA was mixed with 400 µL competent cells and incubated further for 2 hours before plating on selective agar plates which were incubated overnight at 37 °C.

For interaction experiments on solid media, PDA (Potato dextrose agar, Carl Roth) plates were prepared following the supplier’s instruction and plates were dried in a laminar flow cabinet for 1 hour to avoid excessive expansion of *B. subtilis* colonies.

### Population dynamics

Overnight cultures of MB9_B1 mKate2, MB9_B6 GFP, and MB9_B1 mKate2 Δ*ppsC* were adjusted to 0.1 at OD_600_, and mixed in ratios of 100:1, 1:1, 1:10, and 1:100 (MB9_B1/MB9_B1 Δ*ppsC* and MB9_B6 respectively), and used in the experimental setups outlined below.

### Population dynamics in liquid culture

Spores of *F. oxysporum* were harvested from a 5-day grown culture cultivated in 50 ml PDB with shaking at room temperature and filtered through miracloth to remove remaining mycelia fragments before washing in sterile MQ water. The spore solution was diluted 500× followed by 2-fold serial dilution in PDB in a 96-well microtiter plate and 11 µL of the adjusted single strains and mixes (1:1 and 1:10) were subsequently added into the microtiter plate to a final volume of 111 µL.

Growth of the bacterial populations in the presence or absence of *Fusarium* spores were followed using a microplate reader (BioTek Synergy HTX Multi-mode Microplate Reader). The plate was incubated at 28 °C with shaking and measurements of OD_600_, green fluorescence for MB9_B6, and red fluorescence for MB9_B1 were taken every 15 minutes for 70 hours (Optics position: Bottom, GFP: ex: 485/20 em: 528/20, Gain: 50, mKate2: Ex: 590/20, Em: 635/20, Gain 50).

### Population dynamics on solid media

5 µL of the single strains and mixed co-cultures (1:1, 1:10, and 1:100) were spotted approximately 3 cm from the center of PDA medium containing plates and allowed to dry before spotting of 5 µL of *F. oxysporum* spore suspension in the center of the plates. The samples were incubated at room temperature. Growth and inhibition patterns were imaged every hour for 7 days using a ReShape imaging system (ReShape biotech, Denmark) and annotated for when *Fusarium* reached the edge and outer center of the bacterial colonies. For fluorescence imaging, long pass emission filters at 530 and 630 nm and excitation at 467/5 nm and 527/5 nm for green and red fluorescence, respectively. Fluorescence images were processed using ImageJ, briefly, image stacks were converted to 8-bit and colony ROIs were selected from the normal light image by auto-thresholding using Otsu’s algorithm. Each fluorescence channel was measured for total area of the colony and the raw integrated density, which is a sum of pixel values in the selected area.

## Results

### Plipastatin is protective in a well-mixed liquid culture

The inhibitory effect of the *B. subtilis* and its secondary metabolite plipastatin on *Fusarium* is well documented using single bacterial cultures on agar medium (Kiesewalter *et al*. 2021; Kjeldgaard *et al*. 2022); however, its role within multi strain systems is less well understood. To dissect how plipastatin influences the dynamics between different *B. subtilis* strains, we established a two-species co-culture system consisting of a plipastatin producer (MB9_B1) and a co-isolated non-producer isolate (MB9_B6) in the presence of germinating *F. oxysporum* spores. MB9_B1 and MB9_B6 have earlier been studied and shown to be close relatives (ANI of 99.95 for the four housekeeping genes *gyrA, recA, dnaJ,* and *rpoB)* and non-antagonistic towards each other, with MB9_B6 featuring a point-nonsense mutation in *ppsB* resulting in inability to produce plipastatin (Kiesewalter *et al*. 2021). As a control, a plipastatin deficient mutant of MB9_B1 (MB9_B1 Δ*ppsC*) was also co-cultivated in the presence of MB9_B6 to detect the influence of any other genetic differences between the two isolates.

Our liquid culture approach assayed survival and growth of the mono- and co-cultures of MB9_B1 or MB9_B1 Δ*ppsC* and MB9_B6 against serial 2-fold dilutions of a freshly prepared *F. oxysporum* spore suspension ranging from 2-fold dilution to a 128-fold dilution. Co-culture populations were further split into ratios of 100:1, 1:1, and 1:100 of MB9_B1 and MB9_B6, respectively, to assay possible density dependent protection.

In liquid media, the *Fusarium* challenge often resulted in a collapse of the bacterial culture inferred from the loss of fluorescence signal (**Fig. 1, grey lines**) or sharply downwards trajectory before the end-point compared with the control (**Fig. 1, orange lines**). The isolate in lower concentration was undetectable from the background in 100:1 and 1:100 ratios, and these were excluded from further analysis (**Fig. S1** and **S2**).

**Figure 1.**
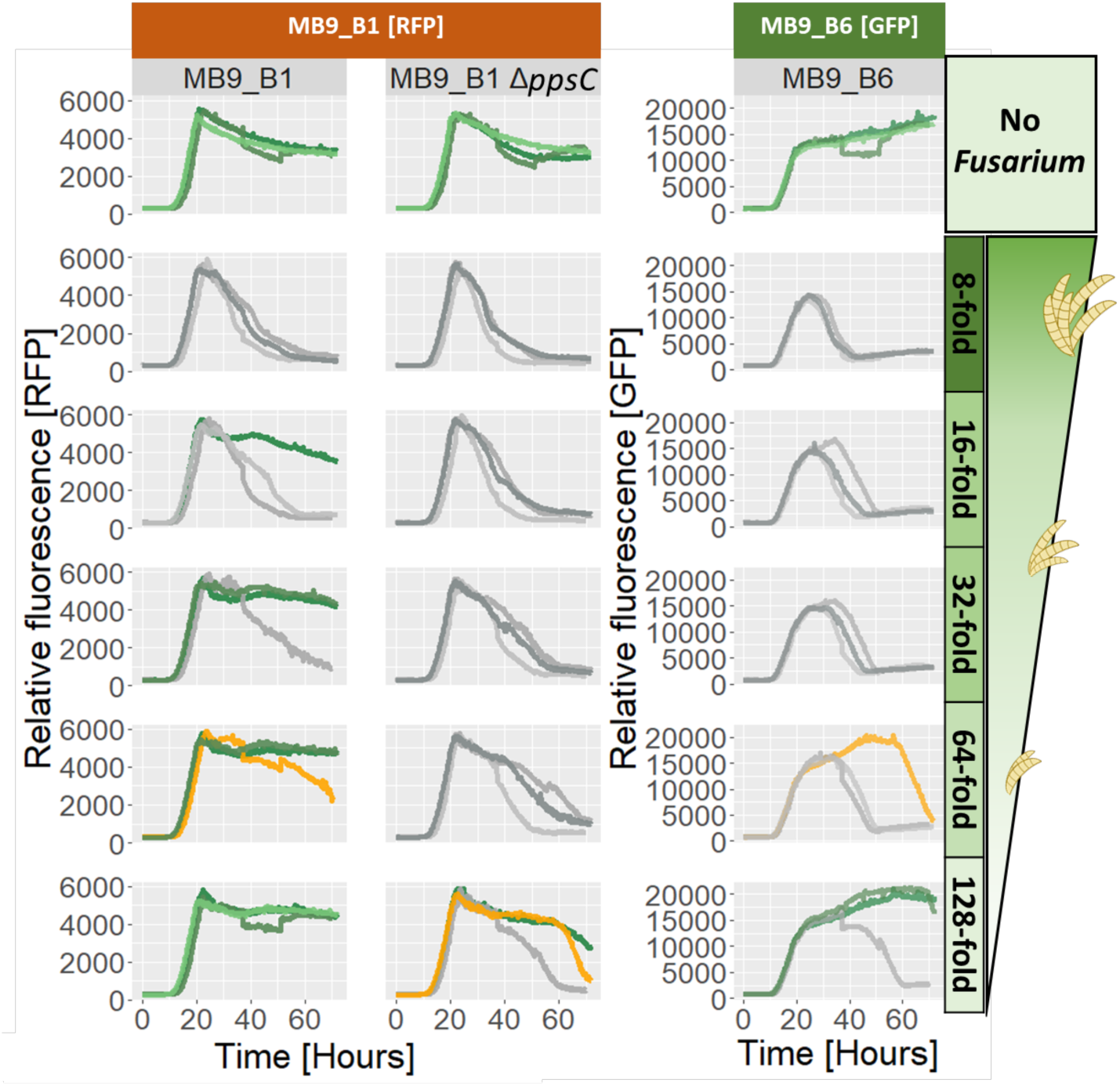
Survival of mono-cultures when challenged with different concentrations of *F. oxysporum* spores in liquid cultures at 28 °C over the course of 70 hours. MB9_B1 and Δ*ppsC* mutant was followed by red fluorescence while MB9_B6 was followed by green fluorescence. The bar to the right denotes the *Fusarium* concentration, with no-spores at the top, followed by the lowest spore dilution and increasing dilutions and thus lower concentration of spores towards the bottom. Surviving populations are colored in green, whereas populations that crashed judging by fluorescence signal are grey, and populations with sharp declines in the late stage are orange. Three replicates of each mono-culture were followed.

All three mono-cultures exhibited similar population crashes at high *Fusarium* loads of 2-, 4-, and 8- fold (2- and 4-fold included in supplementary figures **Fig. S3, S4,** and **S5**), but differentiated at the 16-fold dilution and lower spore concentrations (**Fig 1, green lines**). MB9_B1 populations survived with 1/3 populations at 16-fold, 2/3 at 32- and 64-fold and all 3/3 at 128-fold (**Fig. 1, left panel**) whereas MB9_B6 only featured 2/3 survivors at 128-fold (**Fig. 1, right panel**) and MB9_B1 Δ*ppsC* featured two sharply decreasing populations even at 128-fold dilution of *Fusarium* spores (**Fig. 1, middle panel**).

The differences in survival indicated certain degree of plipastatin-mediated protection against germinating *F. oxysporum* spores in our setup. Next, we determined whether this effect was shared or privatized by the producing strain MB9_B1.

Co-cultures were generally less robust against *Fusarium* invasion, likely due to halving of the producer density. In the 1:1 mixes, MB9_B1 did not survive at the 16-fold dilution or higher spore concentrations, and featured 1/6, 3/6, and 5/6 survivors at 32-, 64-, and 128-fold respectively (**Fig. 2, left panel**) while MB9_B1 Δ*ppsC* only showed 1/3 surviving populations at 128-fold spore dilution (**Fig. 2, second panel from the left**). Plipastatin producers were beneficial towards non-producers, as MB9_B6 featured increased survival in the 1:1 co-culture with MB9_B1 with 1/6, 3/6, and 5/6 surviving populations respectively at 32-, 64- and 128-fold dilutions (**Fig. 2, third panel from the left**) versus 1/3 at the 128-fold dilution when MB9_B6 was co-cultured 1:1 with MB9_B1 Δ*ppsC* (**Fig. 2, right panel**).

**Figure 2.**
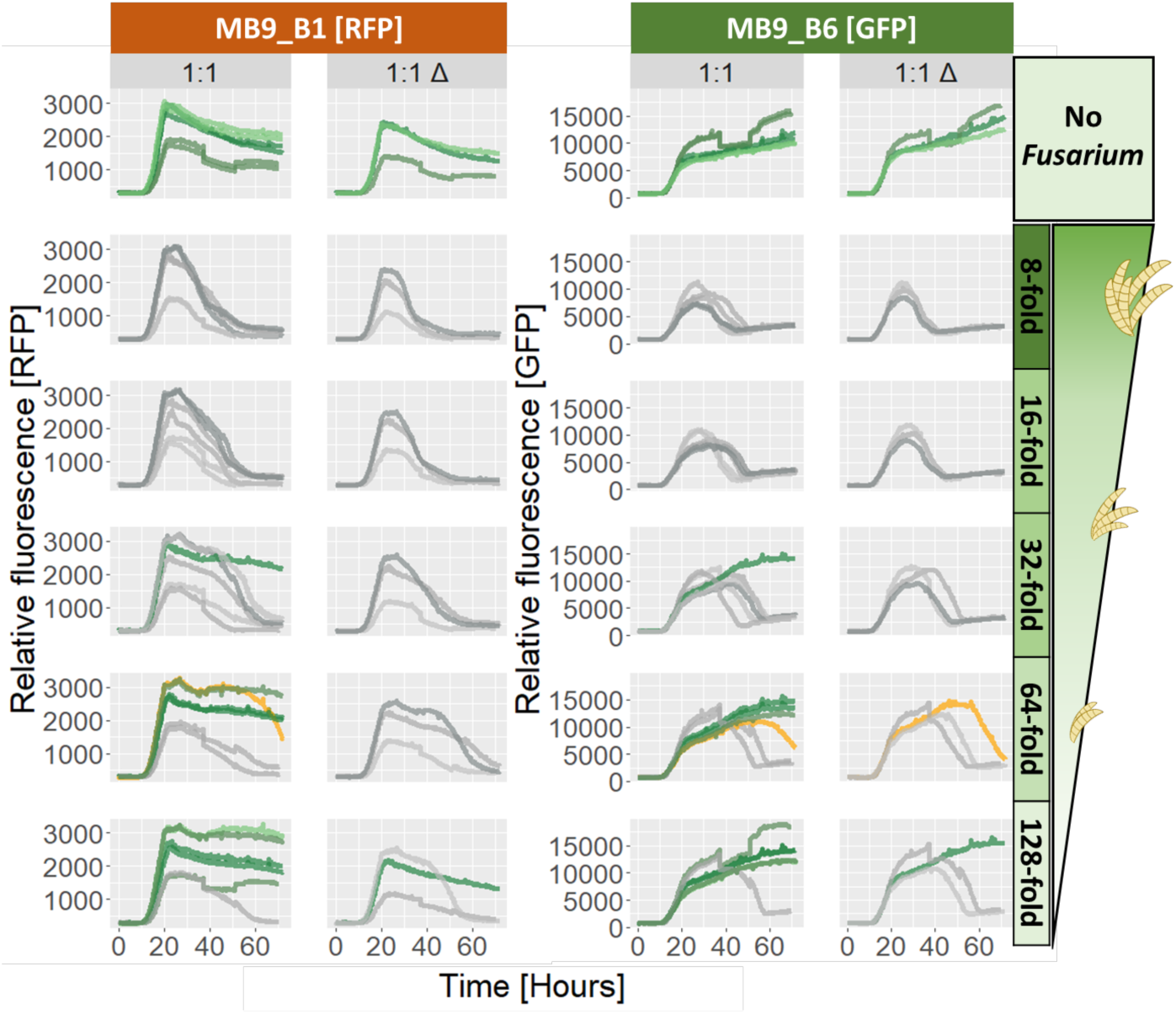
Survival of co-cultures when challenged with different concentrations of *F. oxysporum* spores in liquid cultures at 28 °C over the course of 70 hours. Co-cultures with wild-type MB9_B1 are marked 1:1, while co-cultures using the non-producer mutant MB9_B1 Δ*ppsC* are marked 1:1 Δ. MB9_B1 and Δ*ppsC* mutant in the co-culture was followed by red fluorescence while MB9_B6 was followed by green fluorescence. The bar to the right denotes the *Fusarium* concentration, with no-spores at the top, followed by the lowest spore dilution and increasing dilutions and thus lower concentration of spores towards the bottom. Surviving populations are colored in green, whereas populations that crashed judging by fluorescence signal are grey, and populations with sharp declines in the late stage are orange. 1:1 culture was performed in 6 replicates while 1:1 Δ was performed in 3 replicates.

As co-cultures proved somewhat beneficial to the non-producer, we further inspected the growth of each species without *Fusarium* challenge to determine whether the non-producer had a fitness benefit over the producer. Here, we noted only minor differences in growth characteristics of the monoculture although with MB9_B6 performing best followed by MB9_B1 Δ*ppsC* and close after MB9_B1 (**Fig. 3A**). However, in 1:1 co-culture, MB9_B1 wild-type featured higher fluorescence levels compared to MB9_B1 Δ*ppsC* (**Fig. 3B, middle panel)**, while MB9_B6 fared slightly better in co-culture with the plipastatin non-producer (**Fig. 3B, right panel**), which could indicate a role of plipastatin in intraspecific competition or epistatic effects of plipastatin deletion in our isolate. Overall, our data did not display fitness difference due to the lack of plipastatin production within the tested timeframe and condition.

**Figure 3.**
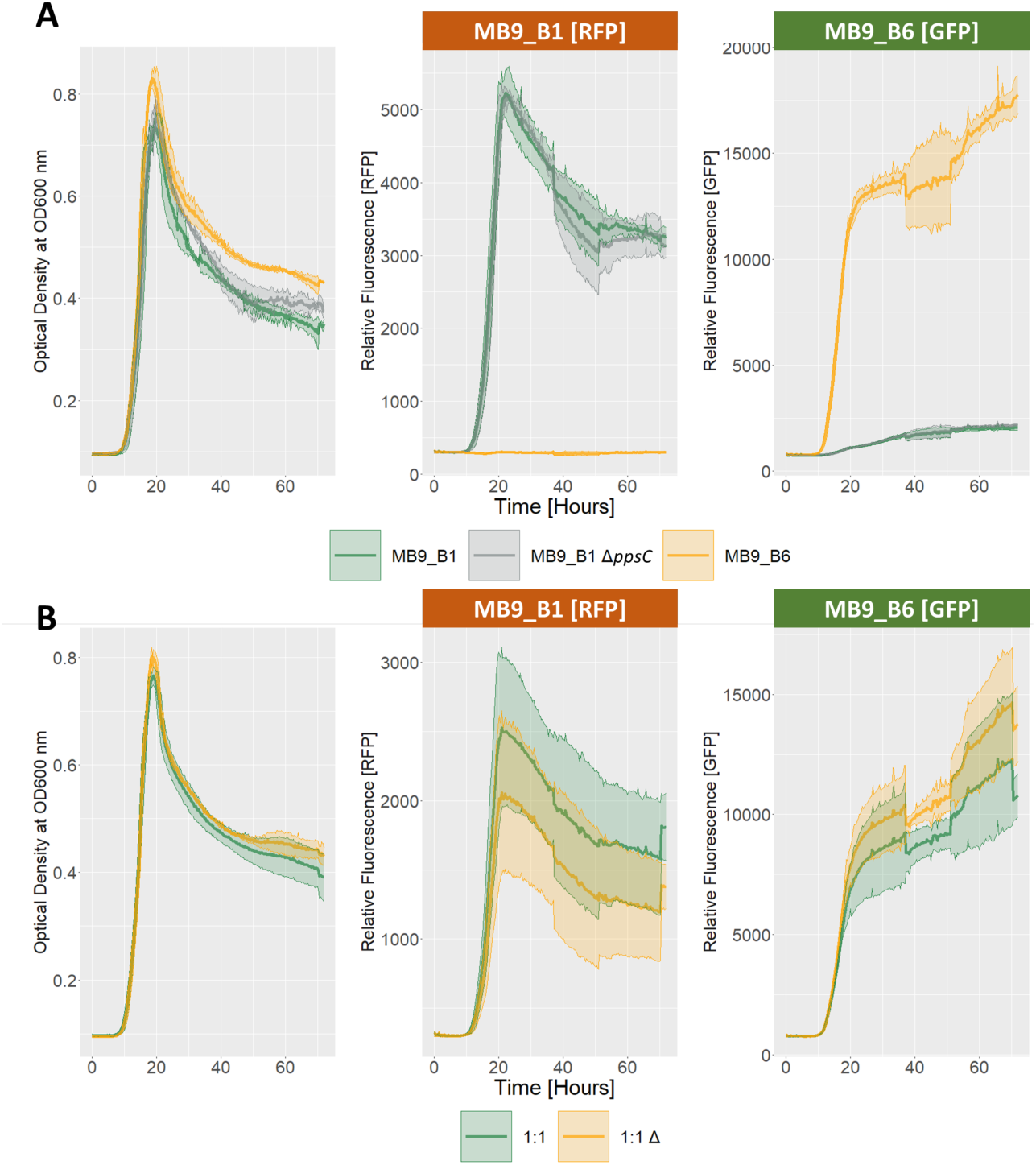
Growth of mono- and co-cultures in the absence of *Fusarium* challenge, incubated at 28 °C for 70 hours and measured for optical density at 600 nm and fluorescence signal from RFP and GFP. (A) Mono-cultures of MB9_B1 (green), MB9_B1 Δ*ppsC* (grey), and MB9_B6 (orange), left panel showing OD_600_, middle panel showing red fluorescence as a proxy for MB9_B1 and Δ*ppsC* mutant, right panel showing green fluorescence as a proxy for MB9_B6. Line shows the mean of 3 replicates with ribbon denoting standard deviation. (B) Co-cultures of MB9_B1 and MB9_B6 (green) and MB9_B1 Δ*ppsC* and MB9_B6 (orange). 1:1 culture was performed in 6 replicates while 1:1 Δ was performed in 3 replicates, line shows the mean of these replicates and ribbon represents the standard deviation.

### 3.2 Protection by the shared plipastatin depends on a structured environment

In nature, bacteria are often found in spatially structured environments, including biofilms, which lacks the homogeneous milieu of a well-mixed liquid culture, which might favor cheaters due to constant mixing. Therefore, we next tested whether plipastatin was also protective towards a non-producer on solid agar plates. On solid agar medium, bacterial colonies and *Fusarium* spores were spaced 3 cm’s apart and left at room temperature to grow, expand, and interact with each other and timed for when *Fusarium* hyphae reached the edge of the colony (**Fig. 4A**) and the colony center (**Fig. 4B**).

**Figure 4.**
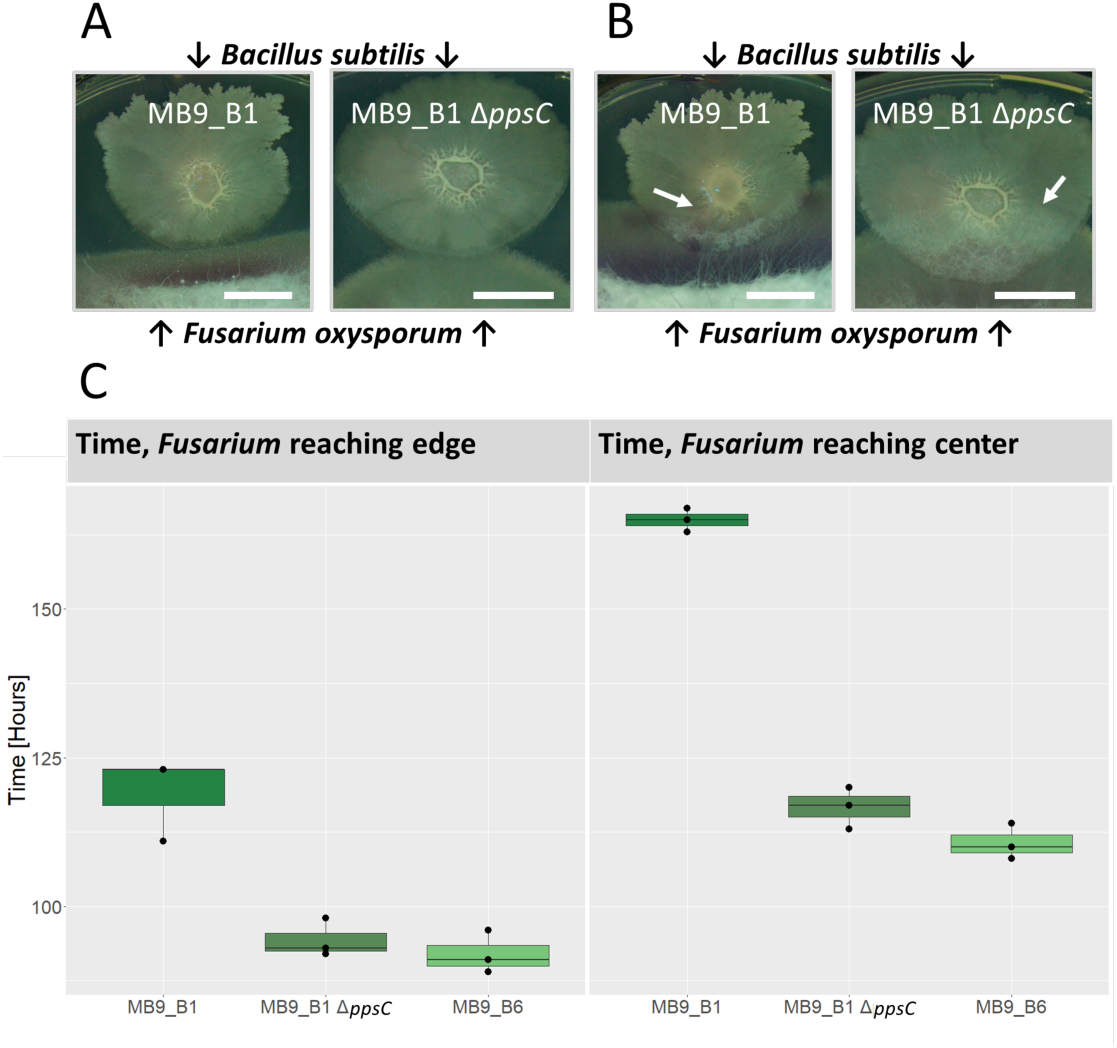
Interactions between *B. subtilis* mono-cultures and *Fusarium* on solid media PDA, grown for 7 days at room temperature. Representative images of when *F. oxysporum* was determined to meet *B. subtilis* colonies at the edge (A) and center (B) with white arrows pointing out the less visible *Fusarium* front when invading into the center. Scale bars indicate 1 cm. (C) Boxplot displaying the timing of *F. oxysporum* encountering edge (left panel) and center (right panel) of mono-culture colonies of MB9_B1, MB9_B1 Δ*ppsC*, and MB9_B6 with three replicates of each.

Generally, given enough time, *F. oxysporum* was able to overgrow *B. subtilis* colonies regardless of plipastatin production, but with considerable difference in the time points when hyphae reached the edge and center of the colony. For monoculture colonies, a clear inhibitory zone was observed for plipastatin producers with accompanying delay in *Fusarium* invasion when compared to non-producers, taking 25 hours longer before *Fusarium* reached the edge and 51 hours before expanded until the center of the colony (**Fig. 4C**) at which time non-producers have already been fully overgrown by *Fusarium*.

Plipastatin production was beneficial in co-cultures with invasion delay depending on the relative abundance of the producer strain, *Fusarium* reached the colony edge of the 1:100 culture first at 90 hours, followed by 1:1 at 100 and finally the 100:1 colony at a 108 hours delay (**Fig. 5A**). In all cases, the delay observed was smaller than that of the producer monoculture, which was likely due to the lower producer density compared with monoculture producer strain, MB9_B1. The importance of plipastatin was further underlined by co-cultures replacing MB9_B1 with its plipastatin non-producer derivative, MB9_B1 Δ*ppsC*, in which no inhibitory zone or delay was observed for *F. oxysporum* (**Fig. 5B**). The time point when *F. oxysporum* reached the center of the bacterial colonies followed a similar trend as detected for the colony edge.

**Figure 5.**
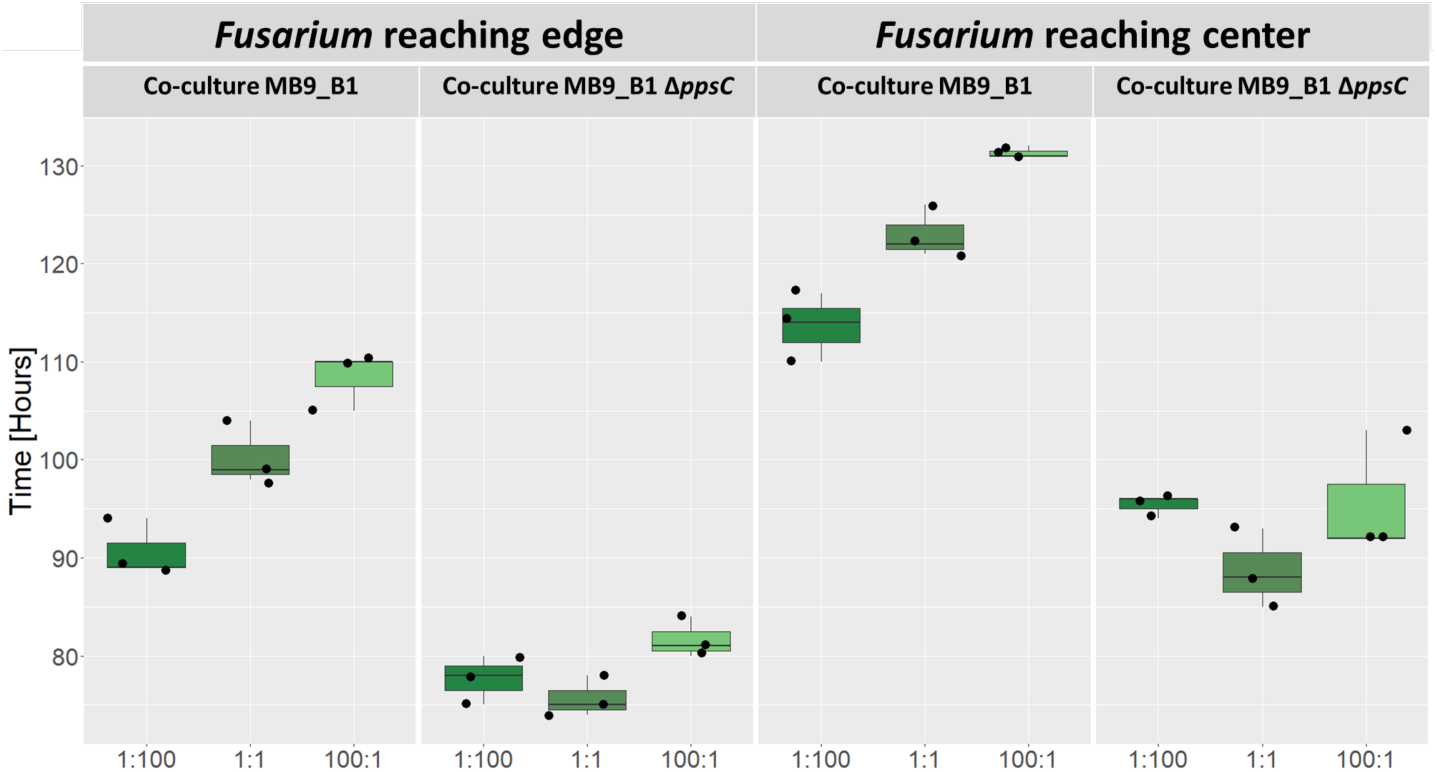
Interactions between *B. subtilis* co-cultures and *Fusarium* on solid media PDA, grown for 7 days at room temperature. Boxplot showing the timing of *F. oxysporum* encountering edge (left top panel) and center (right top panel) of co-cultures of MB9_B1 and MB9_B6 (left sub-panels) and MB9_B1 Δ*ppsC* and MB9_B6 (right sub-panels) at ratios of 1:100, 1:1, and 100:1 of MB9_B1 and MB9_B6 respectively with three replicates of each.

Fluorescence labeling of the strains allowed us to roughly estimate the abundance of producers and non-producer populations in the co-culture colonies. Using the summed pixel values of the colony adjusted by its area for both fluorophores, we could assay the differences between co-cultures without *Fusarium* challenge at day 5 of the experiment as a control. We noted little difference between co-cultures featuring MB9_B1 or MB9_B1 Δ*ppsC* for both red and green fluorescence, suggesting that plipastatin production had little effect on the composition of the co-cultured colonies alone (**Fig. 6)**.

**Figure 6.**
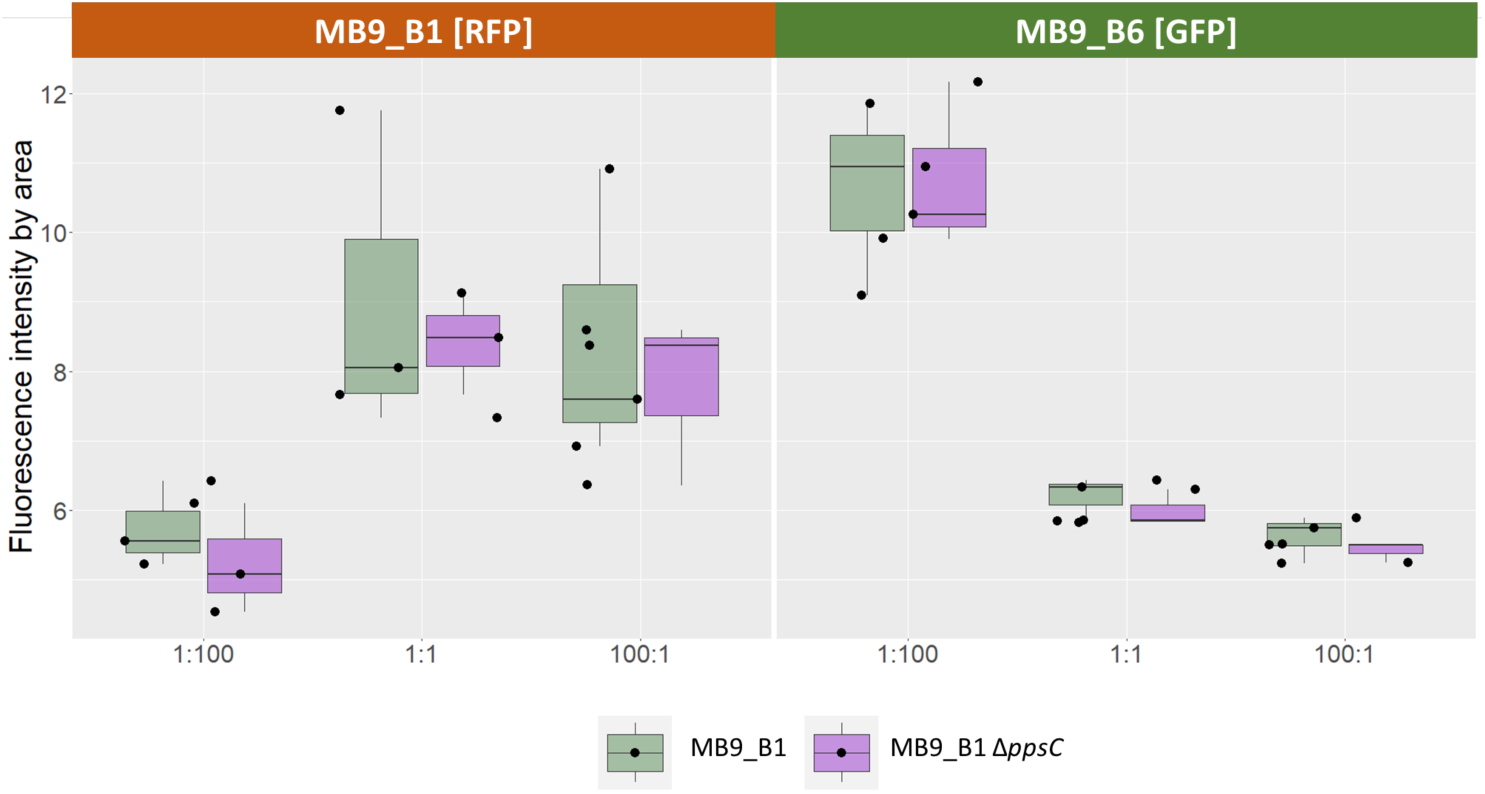
Summed fluorescence intensity divided by the area of colonies for each co-culture without *Fusarium* challenge at day 5 with MB9_B6 at ratios of 1:100, 1:1, and 100:1 of MB9_B1 and MB9_B6 respectively. Red fluorescence of MB9_B1 shown in left panel and green fluorescence of MB9_B6 shown in right panel. Co-cultures with MB9_B1 shown in light green and co-cultures with MB9_B1 Δ*ppsC* shown in purple, boxplots are the result of three replicates.

## Discussion

The efficacy of biological control agents are highly dependent on the context of the environment as the resident microbiome members can interact with the biocontrol strain and its effector molecules. Here, we focused on the population dynamics of bioactive secondary metabolite sharing. Beneficial extracellular metabolite producing organisms are at risk of cheating, which could compromise the long-term survival of a producer strain. By cheating or exploitation, non-producers might gain a fitness advantage over producers through saving metabolic resources and benefitting from the secreted compounds (so called public goods). Evolutionarily, the producer strain is outcompeted, leading to the loss of the beneficial trait in the population (i.e. tragedy of commons). Therefore, producers must ensure cooperation or direct beneficial behavior towards kins that are more likely to reciprocate. Understanding how producers of active metabolites such as plipastatin function in the context of other strains is immensely important for formulating effective biosolutions against plant pathogens like *F. oxysporum*.

Here, we describe that protection against *F. oxysporum* by plipastatin is shared among *B. subtilis* isolates with variable production of this anti-fungal secondary metabolite. Our experiments indicate that protection by plipastatin is dependent on the density of the *B. subtilis* producer strain and also influenced by the level of *Fusarium* spores. Under planktonic conditions, we observed either full inhibition of *F. oxysporum* spore germination and growth or collapse of the bacterial co-culture containing the mixture of plipastatin producer and non-producer strains. In the solid agar media setup, *Fusarium* and *B. subtilis* were spatially separated at the beginning and allowed to meet during growth. Here, presence of plipastatin delayed the fungal invasion, but since *Fusarium* was not completely eradicated, *B. subtilis* colonies were eventually invaded by the fungus regardless of plipastatin producer strain proportion. The different dynamics observed between planktonic and solid surface cultures suggests that plipastatin functions more efficiently as a preventative measure against *Fusarium* establishment rather than during direct competition.

In the planktonic co-cultures, complete and early inhibition of *Fusarium* protected the cohabiting non-producing strain, which raises the question of whether the plipastatin producing strains have fitness a disadvantage compared with the cohabiting non-producers. However, the plipastatin producer MB9_B1 strain had a similar or even increased fitness compared with non-producer mutant derivative when co-cultured with MB9_B6 under the planktonic conditions. Similarly, no significant difference has been observed for the co-cultures that included either the wild-type or Δ*ppsC* MB9_B1 in combination with the natural non-producer strain, MB9_B4. To exclude other factors between MB9_B1 and MB9_B6 that might have played a role in the lack of fitness difference, a co-culture using MB9_B1 and MB9_B1 Δ*ppsC* should be tested in future studies. Further, the timeframe of our experiments lasted up to 3 days in liquid and 7 days on solid media, which might simply be too short period to detect potential fitness differences or changes in the population dynamics. Potentially, a successive experimental setup, in which the co-culture is repeatedly transferred and challenged, could potentially allow to determine the evolutionary stability of plipastatin production in the mixture of plipastatin producer and non-producer strains.

Additionally, other secondary metabolites might influence the stability of microbial consortia, offering another stabilizing mechanism, like division of labor, whereby each partner organism specialize in production of one or a subset of metabolites or matrix components, which has been documented for *B. subtilis* in earlier literature (Gestel, Vlamakis and Kolter 2015; Liu *et al*. 2015; Dragoš *et al*. 2018; Jautzus, van Gestel and Kovács 2022; Yannarell *et al*. 2023). Following expression of plipastatin and other SMs in each isolate during co-culture and when being challenged by *Fusarium* could shed light on potential task distribution, prudent metabolism, and sub-populations that the co-culture might develop.

While we have demonstrated that plipastatin protects kin non-producers in liquid cultures and on solid agar medium, there are still many questions left unanswered regarding the stability of the behavior, plipastatin expression patterns in the producer population, and the role of other secondary metabolites in inhibiting *F. oxysporum*. However, we hope this study will aid in future understanding of how *B. subtilis* functions within a consortium.

## Acknowledgments

This project was supported by the Danish National Research Foundation (DNRF137) for the Center for Microbial Secondary Metabolites

## Author contributions

Rune Overlund Stannius (Conceptualization, Formal analysis, Investigation, Methodology, Visualization, Writing – original draft), Ákos T. Kovács (Conceptualization, Funding acquisition, Supervision, Writing – original draft).

---

**Figure S1.**
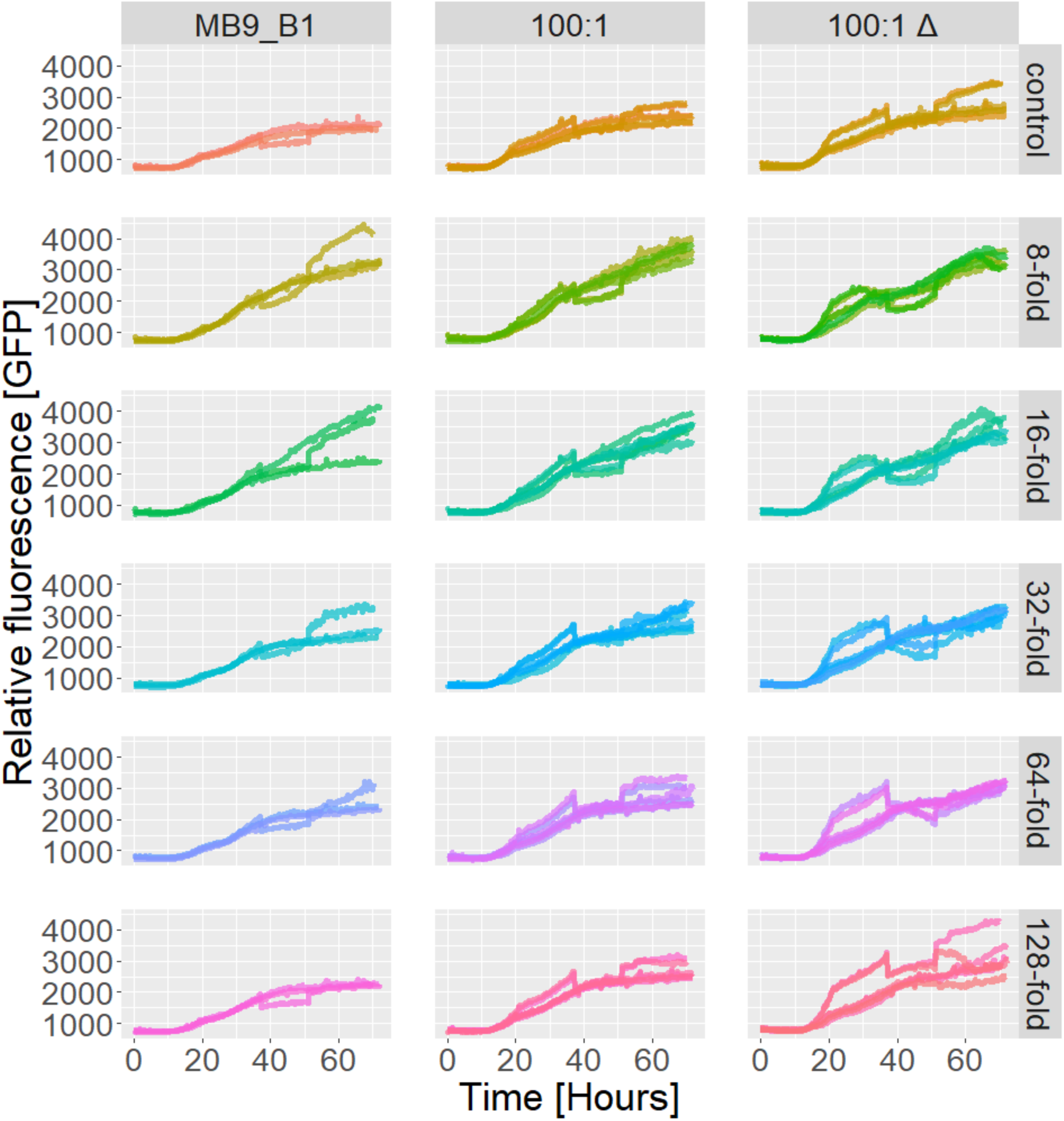
Green fluorescence intensity of the MB9_B1 mono-culture which is not green fluorescent and 100:1 and 100:1 Δ co-cultures showing very low signal of co-cultures compared to the negative mono-culture, and additionally, slightly higher green fluorescence in cultures with *Fusarium* spores.

**Figure S2.**
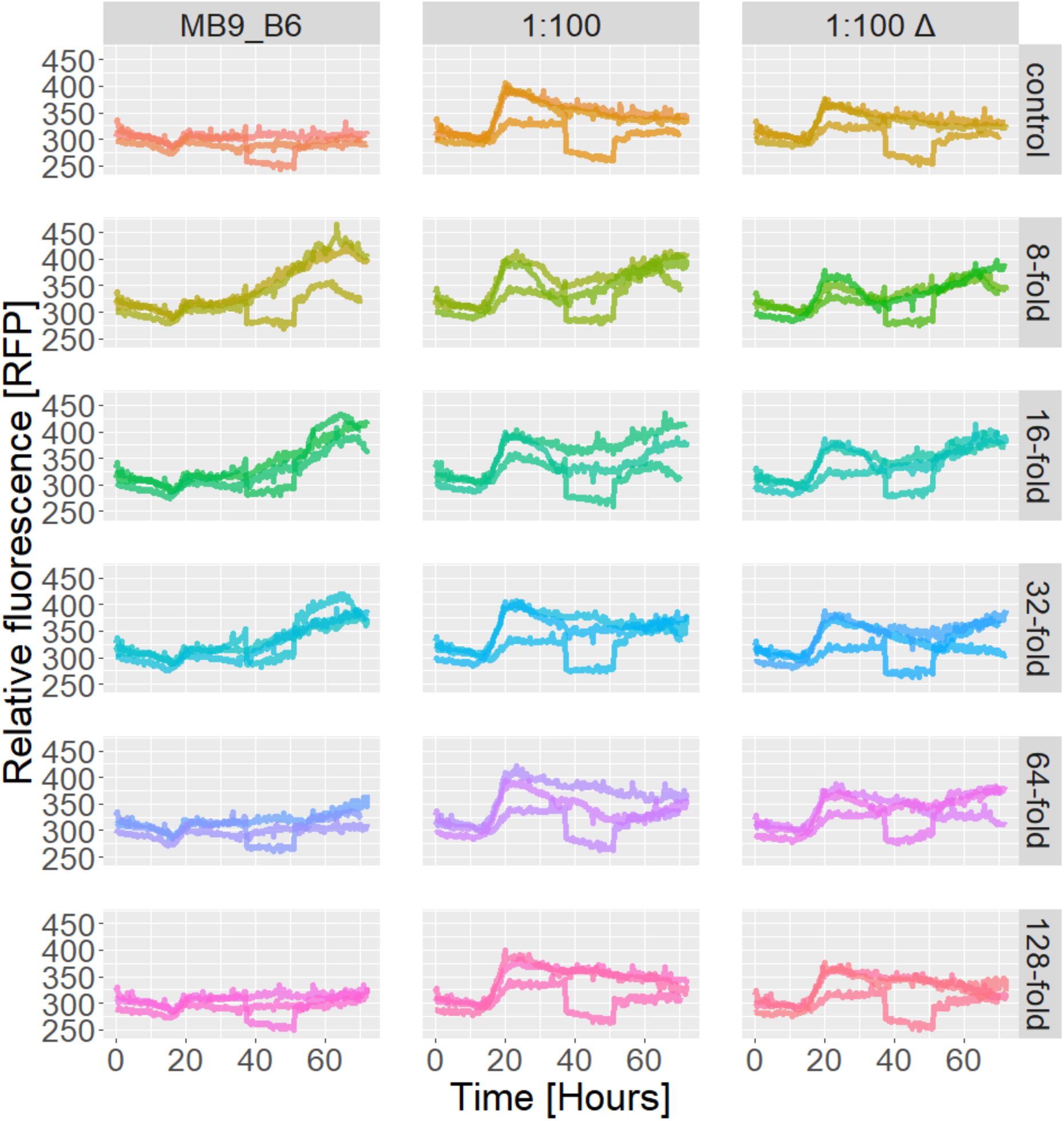
Red fluorescence intensity of the MB9_B6 mono-culture which is not red fluorescent and 100:1 and 100:1 Δ co-cultures showing very low signal of co-cultures compared to the negative mono-culture.

**Figure S3.**
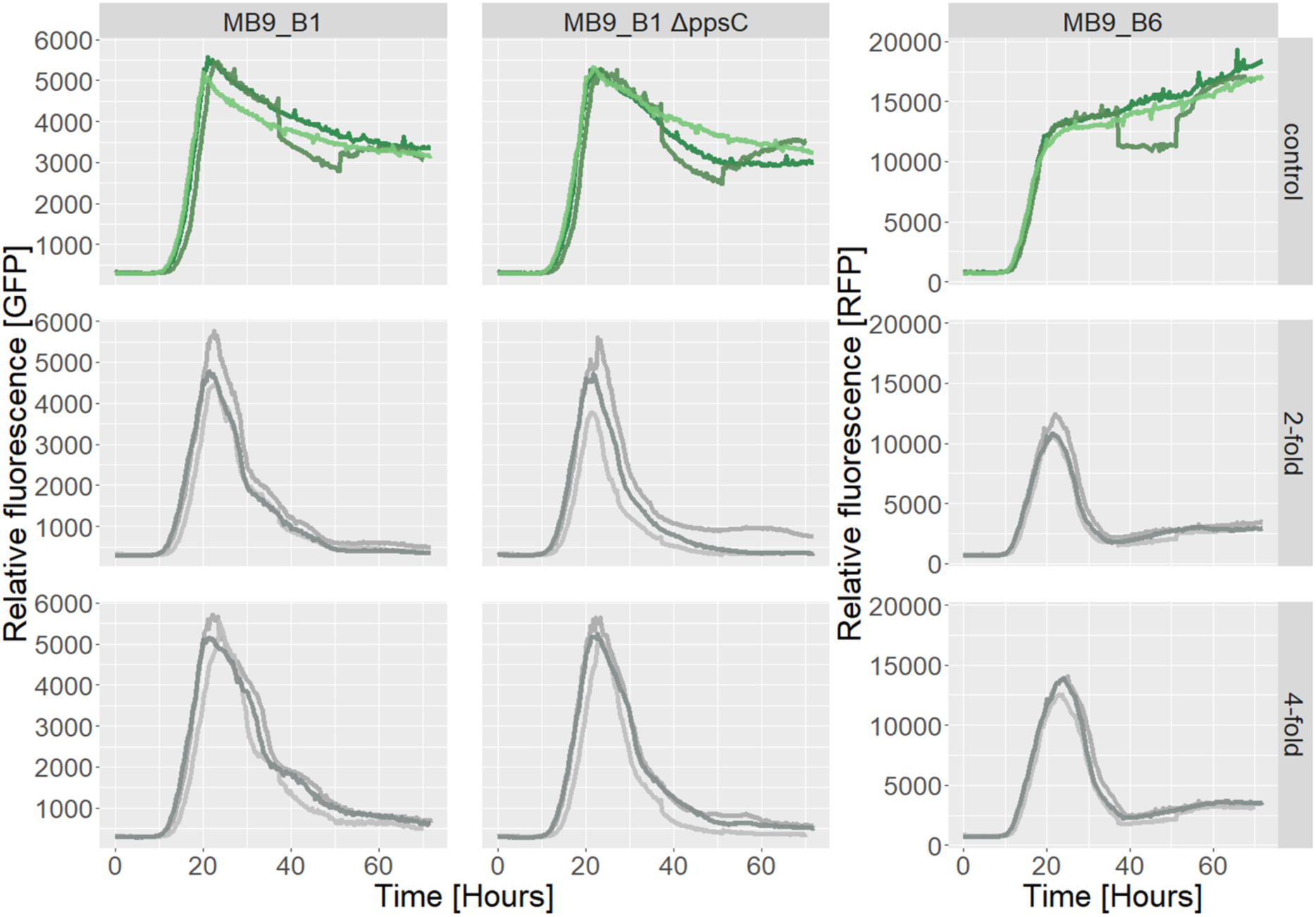
Survival of mono-cultures when challenged with 2-fold and 4-fold dilutions of *F. oxysporum* spores in liquid media at 28 °C over the course of 70 hours. MB9_B1 and Δ*ppsC* mutant was followed by red fluorescence while MB9_B6 was followed by green fluorescence. The bar to the right denotes the *Fusarium* concentration. Surviving populations are colored in green, whereas populations that crashed judging by fluorescence signal are grey. Three replicates of each mono-culture were followed.

**Figure S4.**
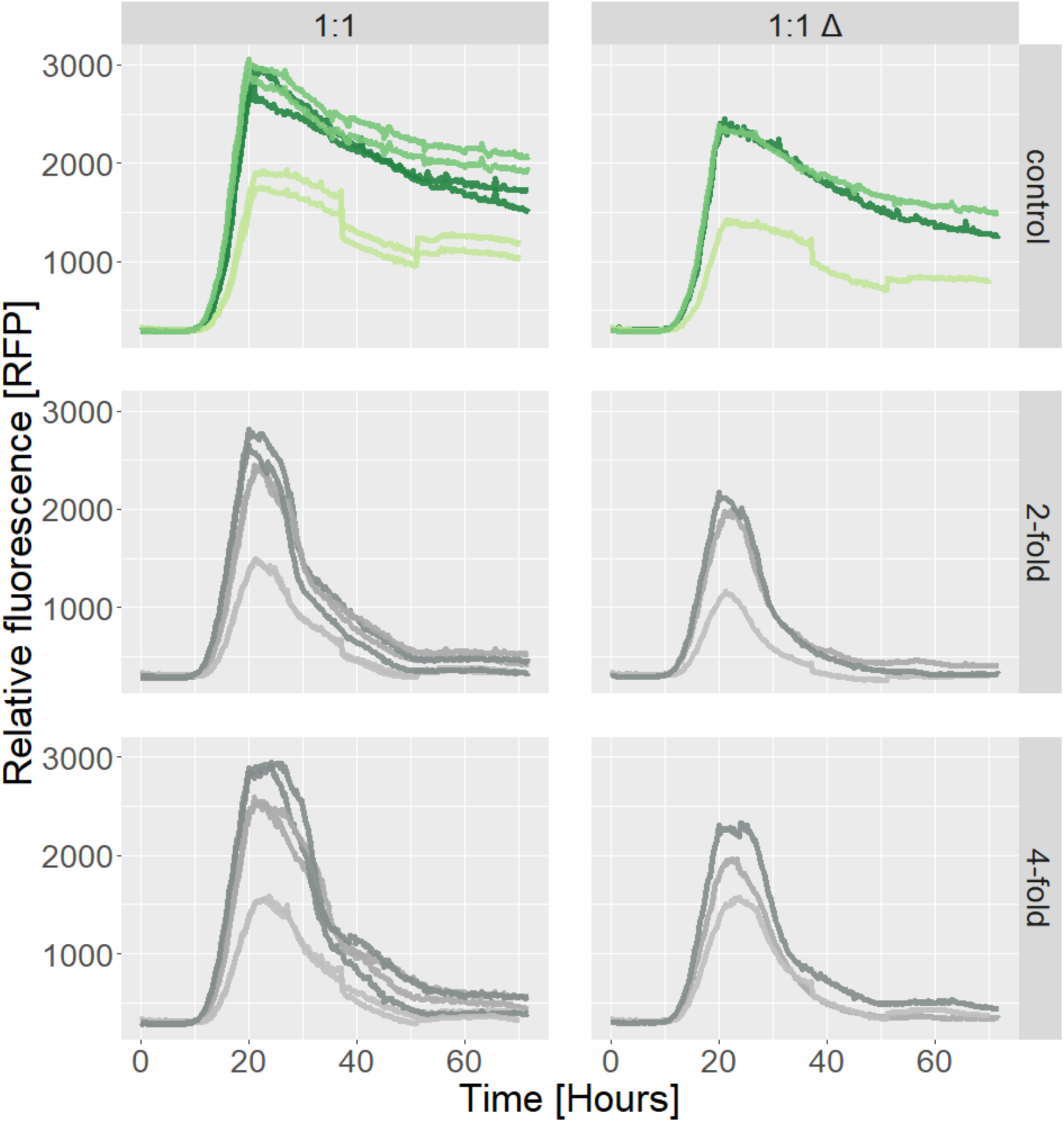
Survival of MB9_B1 by red fluorescence in co-cultures when challenged with 2-fold and 4-fold dilutions of *F. oxysporum* spores in liquid media at 28 °C over the course of 70 hours. Co-cultures with MB9_B1 marked 1:1 (left panel) and co-cultures with MB9_B1 Δ*ppsC* marked 1:1 Δ (right panel). The bar to the right denotes the *Fusarium* concentration. Surviving populations are colored in green, whereas populations that crashed judging by fluorescence signal are grey. Six replicates of the 1:1 co-culture and three replicates of the 1:1 Δ co-culture were followed.

**Figure S5.**
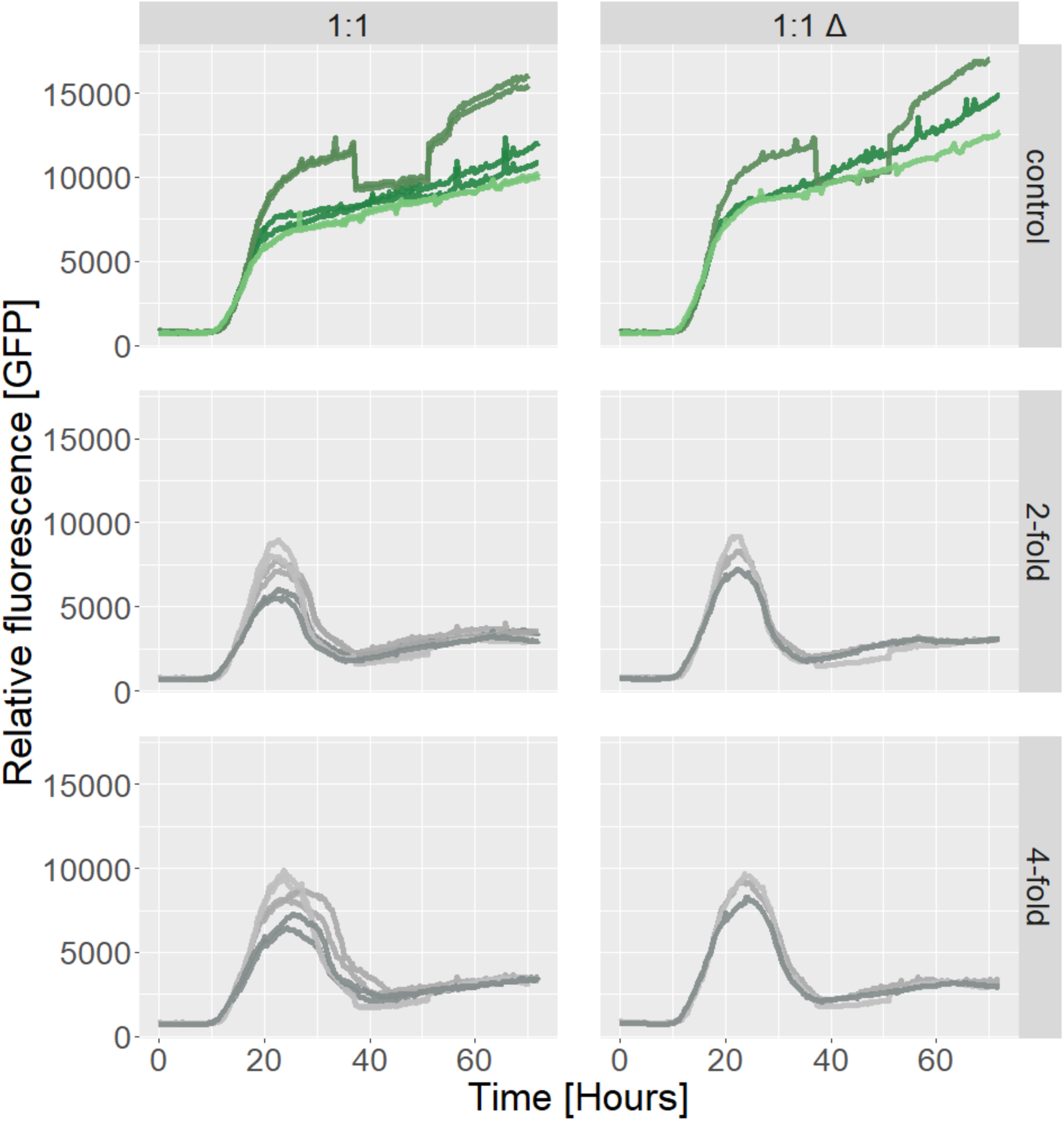
Survival of MB9_B6 by green fluorescence in co-cultures when challenged with 2-fold and 4-fold dilutions of *F. oxysporum* spores in liquid media at 28 °C over the course of 70 hours. Co-cultures with MB9_B1 marked 1:1 (left panel) and co-cultures with MB9_B1 Δ*ppsC* marked 1:1 Δ (right panel). The bar to the right denotes the *Fusarium* concentration. Surviving populations are colored in green, whereas populations that crashed judging by fluorescence signal are grey. Six replicates of the 1:1 co-culture and three replicates of the 1:1 Δ co-culture were followed.

## Notes

### Competing Interest Statement

The authors have declared no competing interest.

